# Pose Estimation of Free-Flying Fruit Flies

**DOI:** 10.1101/2021.01.24.427941

**Authors:** Omri Ben-Dov, Tsevi Beatus

## Abstract

Insect flight is a complex interdisciplinary phenomenon. Understanding its multiple aspects, such as flight control, sensory integration and genetics, often requires the analysis of large amounts of free flight kinematic data. Yet, one of the main bottlenecks in this field is automatically and accurately extracting such data from multi-view videos. Here, we present a model-based method for pose-estimation of free-flying fruit flies from multi-view high-speed videos. To obtain a faithful representation of the fly with minimum free parameters, our method uses a 3D model that mimics two new aspects of wing deformation: a non-fixed wing hinge and a twisting wing surface. The method is demonstrated for free and perturbed flight. Our method does not use prior assumptions on the kinematics apart from the continuity of one wing angle. Hence, this method can be readily adjusted for other insect species.

## I. Introduction

Insect flight is an impressive example of highly maneuverable and robust locomotion [1]. It both challenges our scientific understanding and inspires us to develop tiny bio-mimetic drones [2]. Still, the mechanisms underlying insect flight maneuvers, control and genetics, are elusive and a subject of active study. Modern high-speed cameras and computational tools have greatly advanced insect-flight research. Yet, a significant bottleneck in this field is automatically extracting accurate kinematics from vast amounts of multi-view free-flight videos, where the main challenges are wing deformations and occlusions.

Current tracking methods can be divided into several categories. (1) *Manual tracking*, where a 3D model of the insect is manually fitted to individual frames, is relatively accurate but extremely laborious [3]–[5]. (2) *Landmarks tracking* of feature points on the insect body and wings [6]–[8]. This method might require gluing markers on the insect wings, might suffer from marker occlusion, and often requires manual input. (3) *Deep learning* is a promising method for pose estimation [9], [10], though has not yet been applied to flying insects due to lack of annotated data. (4) *Structured light illumination* has been used to track dragonfly wings and their deformation, but is currently limited to large insects [11]. (5) *Hull reconstruction* methods generate a 3D hull of the insect by tracing the rays from each pixel in each camera view. The hull is segmented into body and wings voxels, from which the insect degrees-of-freedom (DOFs) are extracted [12]–[16]. This approach relies on a generic insect morphology and, hence, can potentially handle a wide range of species. However, its current applications do not handle occlusions very well which might require as many as 8 cameras [16] and often require significant manual input. (6) *Model-based* methods fit a 3D insect model by projecting it onto the camera planes and matching the projections to the data images [17], [18] or by fitting the model to a 3D hull [19]. This approach, first applied for flies in [17], was used in later works (*e.g.* [20]) for analyzing many flight events. Still, obtaining accurate results using this approach requires a 3D model that mimics the insect and its DOFs very faithfully. For example, insect wings are typically not rigid and deform during flight [21], and the wing hinge, connecting the wing to the insect body, is flexible. These deformations cannot be described by modeling the wing as a rigid plate connected at a fixed hinge point.

In this paper, we present a novel work-in-progress model-based algorithm for extracting free-flight kinematics from high-speed multi-view videos of fruit flies. Our 3D model embodies realistic wing deformations using only few additional parameters. This method may alleviate a significant data analysis bottleneck, allowing us to analyze complex phenomena, such as flight control and sensory integration, with high statistical power.

## II. Problem definition

We aim to solve the pose estimation problem for fruit-flies (*Drosophila melanogaster*) in free flight. The input consists of multi-view videos of a fly, and the output is its body and wing kinematics. Body parameters (Fig. 1a) consist of 6 DOFs: 3 translational DOFs and 3 Euler angles (roll, pitch, yaw). The wing parameters are Euler angles that represent wing rotation (Fig. 1b): the stroke angles *ϕ*_*ℓ*_, *ϕ*_*r*_ represents the wing’s forward and backward sweeping motion within the stroke plane; the elevation angles *θ*_*ℓ*_, *θ*_*r*_ describes wing elevation with respect to the stroke plane; and the wing-pitch angles *ψ*_*ℓ*_, *ψ*_*r*_ measures wing rotation around its leading edge. Thus, the minimal kinematic description consists of 12 DOFs.

**Fig. 1.**
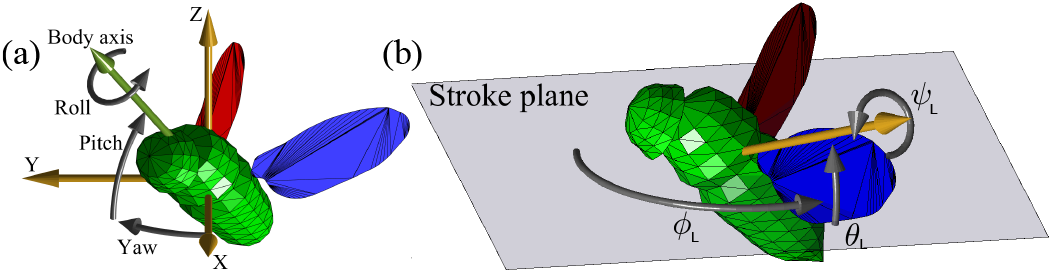
Basic 12 DOF model parameters. **(a)** Body 6 DOF describing its position and orientation. **(b)** Each wing is described by 3 Euler angles relative to the stroke plane: stroke (*ϕ*), elevation (*θ*) and wing pitch (*ψ*). The annotations are for the left wing.

## III. Method

### A. Experimental setup

The experimental setup (Fig. 2) consists of 3 orthogonal high-speed cameras (Phantom v2012, Vision Research), operating at a rate of up to 22,000 frames/sec and 1280×800 pixel resolution. The cameras are back-lit by IR LEDs and tilted upwards by ~36° to reduce wing-wing and body-wing occlusions with respect to the standard Cartesian camera configuration. The volume mutually seen by the cameras is ~5×5×5 cm^3^, located at the center of a custom-made 3D-printed cage. The camera system is calibrated [22], allowing us to convert between 3D world-points and 2D image-points. 10-30 female flies (2-5 days old) are placed in the cage and recorded as they fly through the filming volume. To study insect flight control, we exert mechanical perturbations to the flies by gluing a tiny magnet to the back of each fly and using a magnetic pulse to rotate it in mid-air [14], [15], [23].

**Fig. 2.**
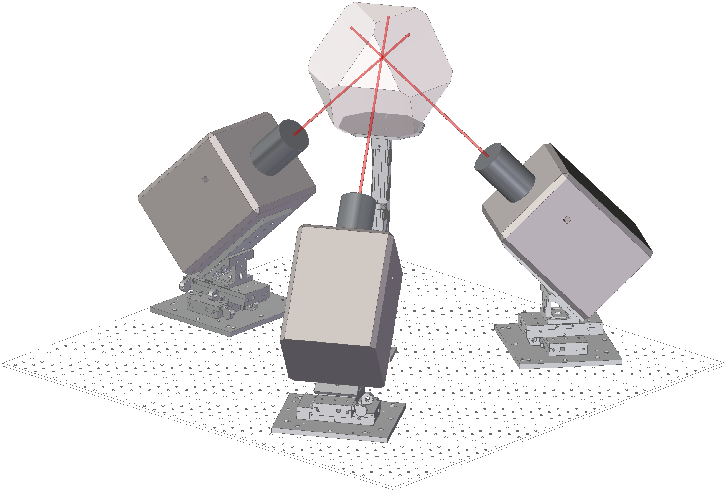
Experimental setup. Three orthogonal high-speed cameras focused on a transparent chamber. The non-Cartesian setup reduces wing occlusions.

### B. Background subtraction

Back-lighting makes the fly pixels darker than the background. Thus, the background is computed by taking the pixel-wise maximum across two frames: the first and the last video frames. To obtain a binary mask from each frame, we first subtract its background, use the transformation *p* → 1 − (1 − *p*)^6^ to deal with wing transparency and apply a binary threshold.

### C. Generative model

Our model for the fly’s body is based on [17] with slight rescaling and a modified head pose. The wing model was obtained by imaging a fly’s wing on a microscope and tracing its outline. The accuracy of model-based pose estimation strongly depends on how well the model and its DOFs mimic the target object. We found that using the 12 DOF description (Fig. 1) leads to significant tracking inaccuracies, because this model does not include two important geometrical features of the fly (Fig. 3). First, due to the flexibility of the wing base, the wing hinge cannot be accurately described as a single point (Fig. 3a). In our model, this feature is described by allowing the two wing hinges to translate symmetrically with respect to the body, which requires 3 additional kinematic parameters: *δx*, *δy* and *δz* hinge translations in the body frame of reference. Allowing asymmetric hinge translation (6 parameters) favored motion of the wing hinges over the wing angles, which hindered the optimization. Second, the wing surface deforms due to the interplay between aerodynamic, inertial and elastic forces acting on the wing [24]. Although these deformations are small, they cannot be captured by a rigid wing model, which introduces sizeable tracking errors, especially during wing pronation and supination (Fig. 3b). In our model, wing deformation is described by a single parameter per wing: *α*_*ℓ*_, *α*_*r*_. As observed experimentally, wing deformation is largest near its base and decreases towards the wing tip [21]. Each *α* parameter quantifies twist per unit length; twist increases linearly from the wing tip (no twist) to the wing base (maximum twist). The model wing is twisted only at the bottom half below its center-line (Fig. 3c).

**Fig. 3.**
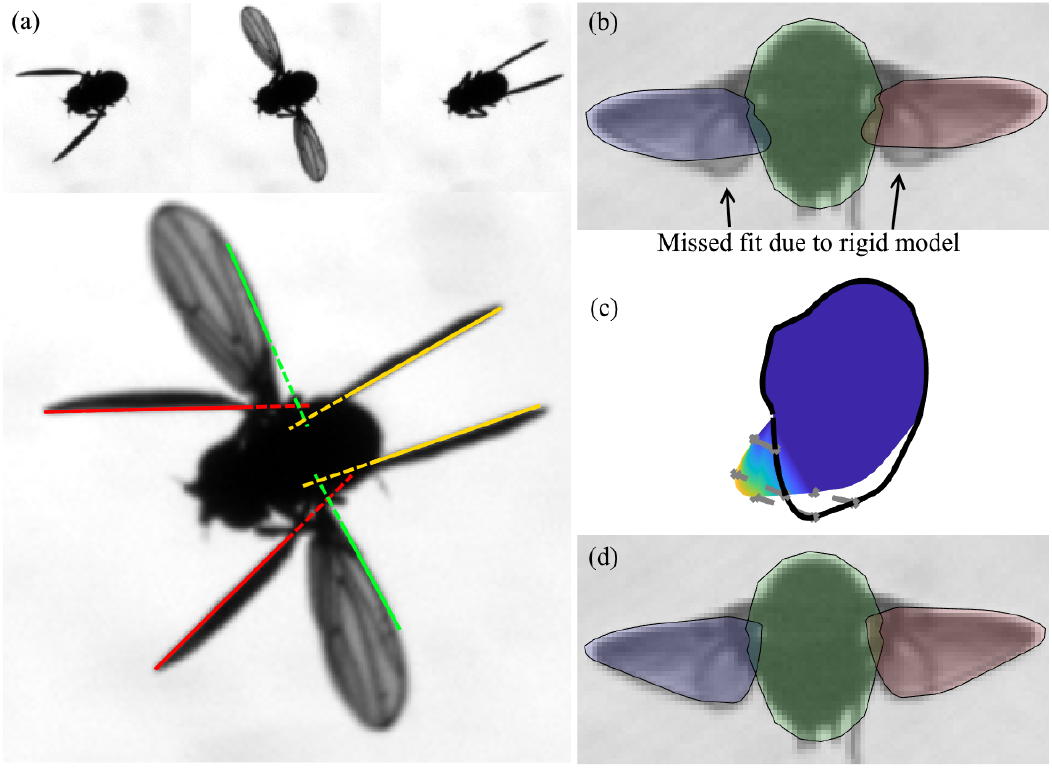
Wing deformations. **(a)** Top: 3 frames from different phases of a single wing beat. Bottom: Superimposing the 3 frames shows that the wing hinge is effectively not fixed during the stroke. The solid lines marking the leading edge of the wing, do not intersect at a single point (dashed lines). **(b)** An unsuccessful fitting attempt using a rigid wing on a frame with twisted wing during supination. **(c)** Wing deformation used in our 3D model. Color represents deformation level and the black line shows the rigid wing outline. **(d)** A successful fit using a flexible wing.

In summary, our model consists of 17 kinematic parameters: the standard 12 DOF, 3 translational offsets of the wing hinge and 2 twist parameters.

### D. Loss function optimization

To quantify the disagreement between the model and a single image, we first project the 3D model onto the corresponding camera plane using the calibrated camera matrix and represent the projection as a 2D polygon. The single-frame loss function is defined as the non-overlapping area (*XOR*) between the model polygon and fly’s binary mask (Fig. 4). The multi-view loss function is a weighted mean of the single-view losses. As tracking the wings is more difficult than tracking the body, we assign greater weight to views that hold more information about the wing pose. The weight of each view, calculated from the initial condition, is proportional to the percentage of wing area unoccluded by the body.

**Fig. 4.**
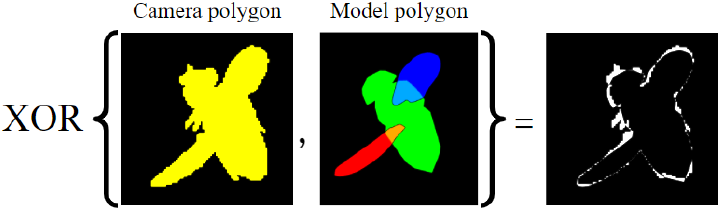
Single-frame loss function. XOR operation on the camera image mask and the projected model.

To evaluate the model parameters at a given timepoint, we minimize the multi-view loss function using a derivative-free interior-point method (*fmincon* in MATLAB). The initial condition for the optimization is the result of the previous frame. The initial condition for the first frame is obtained semi-automatically, where the user applies manual adjustments to automatic optimization results via a graphical user interface. At this step, the user can also determine constant scaling parameters of the model to handle flies of different sizes. The fitted angles were constrained to the entire physiologically possible range for flies. The body roll angle was constrained to ±2° from its initial condition.

We identified that the combination of our loss function and camera configuration leads to degeneracy of the model in certain body and wing poses. As shown in figure 5, two values of the wing-pitch angle *ψ* of the left wing, which differ by ~30°, generate almost identical projections of the model. Consequently, in such cases, optimization might converge to a wrong local minimum. To address this degeneracy, we exploit temporal information by detecting discontinuities in either *ψ* or the loss function. Then, we use multi random start, where we restart the optimization process from 15 random points in parameter space and then re-fit previous ‘suspected’ frames using the same constraints as detailed above.

**Fig. 5.**
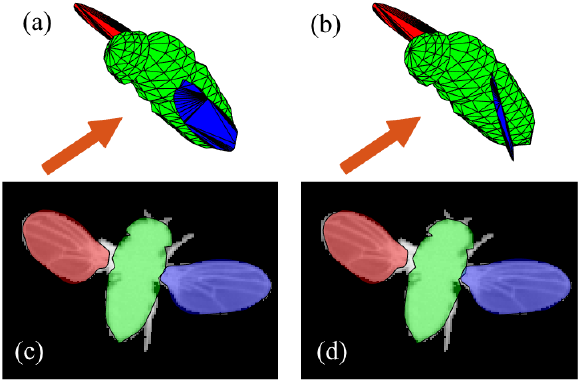
Degeneracy in *ψ*. **(a)-(b)** The 3D model generated by two sets of parameters. The orange arrow shows the direction of the camera taking the images on the bottom. **(c)-(d)** The projection of the corresponding models on the camera plane. The projections are nearly identical.

## IV. Results

### A. Validation

To validate our method, we first tested it on an ensemble of synthetic images generated from the basic 12-DOF model used for optimization. We used previously measured and manually-corrected flight kinematics [14] to generate 36 videos of 100 time points each (a single wing beat). Each video differs by the body yaw angle. Fig. 6 shows a box plot of the resulting errors for each DOF. The fly’s center of mass position was accurate within 10*μ*m (≈0.2 pixel). In the angular parameters, in 98% of the frames the error in all angles was *<*2°.

**Fig. 6.**
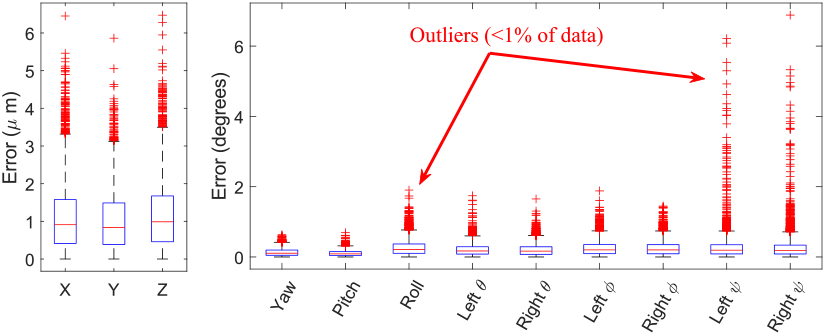
Model validation on synthetic data. Tracking errors box plot. Each box contains 75~ of the data. Whiskers correspond to 99.3~ of the data.

### B. Unperturbed Flight

Fig. 7 and Movie 1 demonstrate pose estimation of a real free-flight sequence. Interestingly, the oscillations in the body pitch angle (Fig. 7b) correspond to the natural periodic pitch motion of the fly: when the wings are in the forward half of the stroke plane (*ϕ<*90) they exert a pitch up torque on the body, and when *ϕ>*90 the wings exert a pitch down torque. Together, these torques result in small, ~2° amplitude pitch oscillations that are clearly seen in both the raw and measured data. Tracking the wing angles (Fig. 7c) shows the typical 8-figure-like trajectory of the wing-tip. The mean loss across the entire movie was 0.1049±0.0068 (mean±standard deviation), better than the loss of fitting the rigid 12 DOF model, which was of 0.1501±0.0204 (Movie 3).

**Fig. 7.**
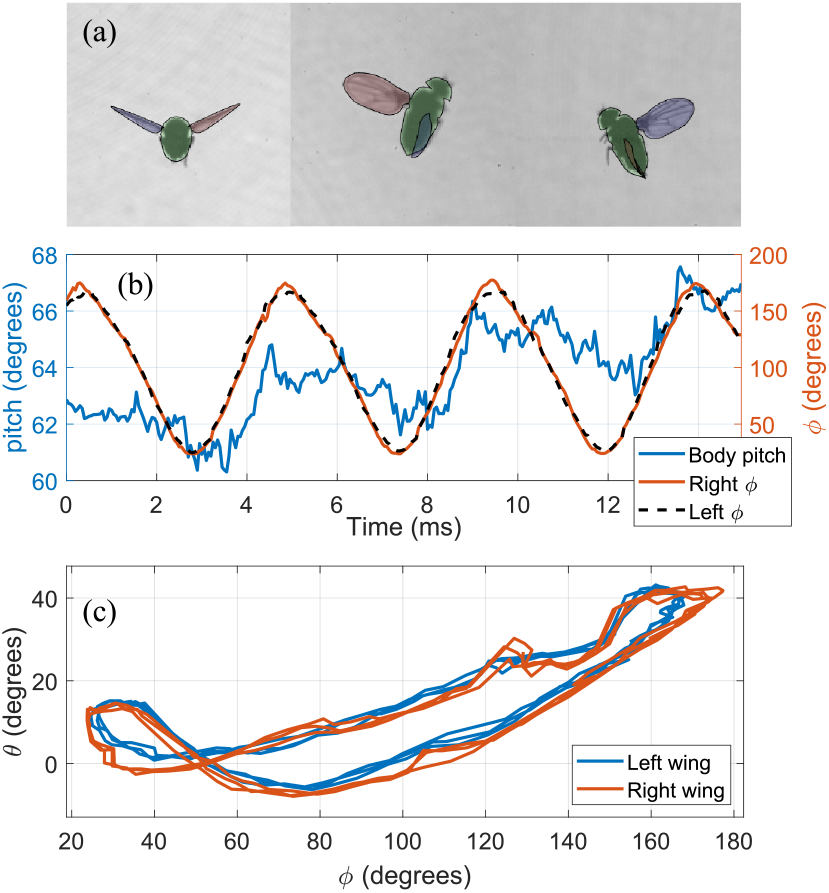
Results on an unperturbed flight event. **(a)** The projection of a fitted 3D model superimposed on the corresponding frames. **(b)** Body pitch and wing *ϕ* **(c)** The path of the wing tip by its elevation (*θ*) and azimuth (*ϕ*)

### C. Roll Perturbation

Fig. 8 and Movie 2 show pose estimation of a real roll correction maneuver in response to a mid-air magnetic perturbation (Section III-A). Here, we modified the 3D model to include the magnetic rod and determined its position manually along with the initial condition. Tracking the body angles (Fig. 8a) shows the fly was rolled to its left by 62° at *t*=17ms after the onset of the perturbation. Body yaw and pitch were also perturbed by −12° and 40°, respectively, because the magnetic torque was not aligned with any body principal axis.

**Fig. 8.**
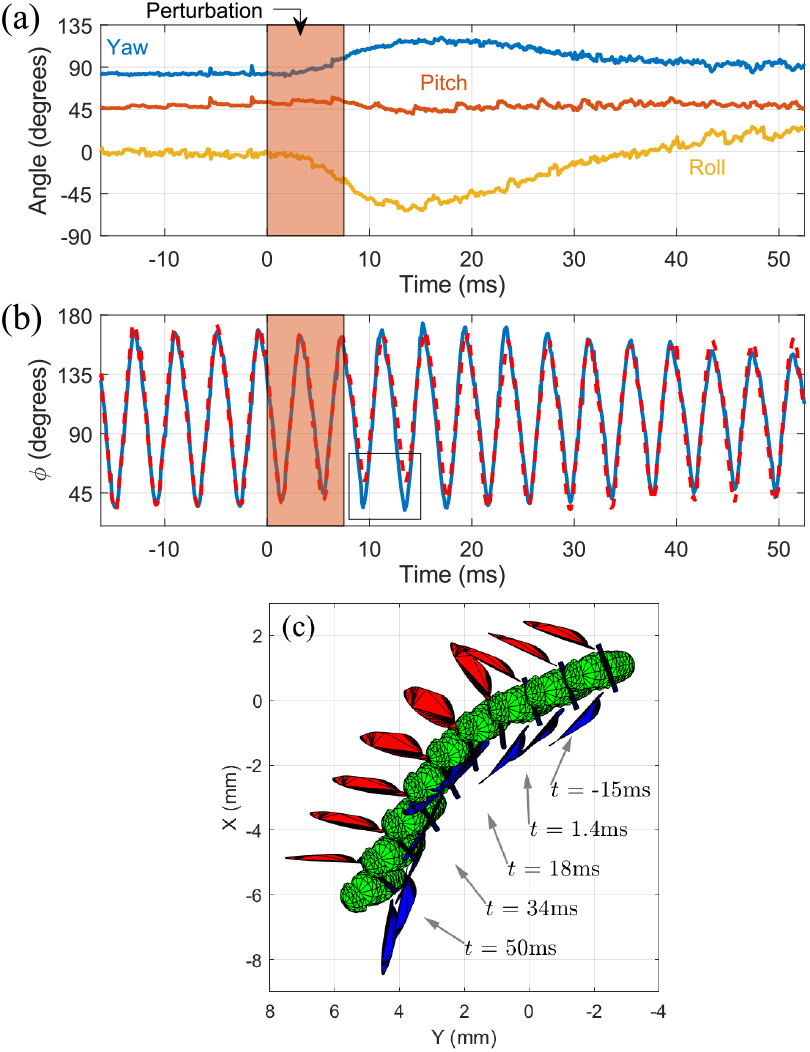
Roll correction. **(a)** Body angles during the maneuver. Magnetic pulse was activated between *t*=0−7.5ms. **(b)** Wings stroke angles. Blue line and red dashed line mark *ϕ*_*ℓ*_, *ϕ*_*r*_ respectively. Rectangle marks the main wing asymmetry during the maneuver. **(c)** Top view of the fitted model shows every two wing beats when the left wing is at supination. Wing stroke asymmetry is clearly visible.

Tracking the wing stroke angles demonstrates the fly’s roll control mechanism [14], where the ‘bottom’ wing (here, left) increases its stroke amplitude and the ‘top’ wing decreases its stroke amplitude. The roll reflex latency was ≈9ms and the perturbation was fully corrected after ~9 wing beats (*t*≈40ms). A characteristic feature of these maneuvers is the residual error in yaw [14], which was 10° in this example.

## V. Conclusion

We presented a pose-estimation algorithm for tracking free-flying fruit flies. The novel features of the model include wing deformation, non-fixed wing-hinge and the addition of magnetic rod for perturbation experiments. Further, our algorithm does not use any prior assumptions on the kinematics, except for the continuity in *ψ* for error detection. Future improvements might involve deep learning using synthetic data and our current results to fully automate the process. Overall, this work-in-progress defines a streamlined data analysis pipeline, that can be easily converted to work with other types of insects.

## Supporting information

Supplemental Movie 1 - unperturbed

Supplemental Movie 2 - roll perturbation

Supplemental Movie 3 - unperturbed 12 DOF fit

## Acknowledgments

We thank Roni Maya, Noam Tsory, Noam Lerner and Bar Karov for assisting in data acquisition. This research was supported by the Azrieli Foundation Faculty Fellowship and by the Alexander Grass Bioengineering Center of the Hebrew University of Jerusalem, Israel.

